# Quantitative comparison of PI(3,5)P_2_ biosensors reveals SnxA is the most sensitive and unbiased

**DOI:** 10.1101/2025.11.10.687688

**Authors:** Tiernan Swayhoover, Claire C. Weckerly, Gerald R. V. Hammond

## Abstract

PI(3,5)P_2_ is an endosomal lipid whose depletion is associated with a variety of pathologies such as neurodegenerative diseases. However, studying this lipid in physiological and disease models has been difficult due to the scarcity of the lipid and the lack of live-cell imaging tools. That is until recently, when a novel PI(3,5)P_2_ biosensor, SnxA, was characterized. Despite the exciting promise of this new sensor, it was still unclear if SnxA unbiasedly reported on PI(3,5)P_2_ levels and how its sensitivity compared to other PI(3,5)P_2_ biosensors. In this work, we addressed these gaps by using a recruitable PIKfyve construct to demonstrate that ectopically generated PI(3,5)P_2_ at mitochondria was sufficient to recruit SnxA. Further, we co-expressed putative PI(3,5)P_2_ biosensors to definitively show that SnxA is more sensitive to PI(3,5)P_2_. We also validated previous results by showing that SnxA depends on PI(3,5)P_2_ for membrane binding, SnxA responds to PI(3,5)P_2_ production at endosomes, and that PI(3,5)P_2_ levels decline quickly when its production is inhibited. Thus, we conclude that SnxA is a robust and sensitive PI(3,5)P_2_ biosensor that facilitates real-time analysis of this key lipid.

## Introduction

Phosphatidylinositol 3,5-bisphosphate (PI(3,5)P_2_ ) is a low-abundance, endosomal lipid first discovered as a metabolite upregulated in yeast exposed to hyperosmotic stress (Dove et al., 1997). PI(3,5)P_2_ is produced by a PIP 5-kinase, PIKfyve in mammals or Fab1 in yeast, that phosphorylates phosphatidylinositol 3-phosphate (PI3P) at the 5-OH position (Cooke et al., 1998; Sbrissa et al., 1999) ). Deletion of this kinase or use of inhibitors such as apilimod results in a phenotype where yeast vacuoles or mammalian endosomal compartments become enlarged; a process called vacuolation (Gary et al., 1998; Ikonomov et al., 2002, 2001; Cai et al., 2013; Jefferies et al., 2008; Rutherford et al., 2006).

PIKfyve exists within a regulatory complex consisting of a homopentamer of the scaffold protein Vac14, one copy of PIKfyve, and one copy of the 5’-phosphatase Fig4, that reverts PI(3,5)P_2_ back to PI3P (Lees et al., 2020). Deletion of PIKfyve is embryonic lethal in mice (Ikonomov et al., 2011) and mutations within the components of the PIKfyve complex have been implicated in neurodegenerative diseases such as Charcot-Marie-Tooth syndrome or amyotrophic lateral sclerosis (ALS) (Chow et al., 2009; Rivero-Ríos and Weisman, 2022; Chow et al., 2007). Although PIKfyve is a druggable therapeutic target (Burke et al., 2022; Ikonomov et al., 2019), the specific role of PI(3,5)P_2_ in these pathologies is still unclear, partly due to a lack of clarity in PI(3,5)P_2_’s subcellular localization.

Our current understanding of PI(3,5)P_2_ localization is based on disease phenotypes, the localization of PIKfyve, and the localization of PI(3,5)P_2_ effector proteins such as WD-repeat proteins interacting with phosphoinositides (WIPI) or membrane channels. Based on this data over the years, PI(3,5)P_2_ has been associated with membranes across the endosomal pathway, including early endosomes, late endosomes, lysosomes, autophagosomes, and the multivesicular body (Cai et al., 2013; Jefferies et al., 2008; Rutherford et al., 2006; Ikonomov et al., 2001; Shisheva et al., 2001; Cabezas et al., 2006; McCartney et al., 2014; Dove et al., 2009; Jeffries et al., 2004; Dong et al., 2010; Banerjee et al., 2019; Leray et al., 2022; Wang et al., 2012). A more recent study that used immunofluorescence to visualize endogenous PIKfyve tagged by CRISPR/Cas9 showed strong co-localization of PIKfyve and the retromer complex, suggesting PI(3,5)P_2_ enrichment on membranes undergoing retrograde trafficking (Giridharan et al., 2022). However, the lack of agreement (and in some cases the presence of direct contradictions) in these collective studies highlights the need for more robust tools to directly monitor PI(3,5)P_2_ .

Genetically encoded lipid biosensors are the gold standard for studying lipids by live cell imaging. Valid biosensors must show selectivity for the lipid of interest *in vitro*, lipid-dependent membrane association, and recruitment to membranes by the lipid alone (Wills et al., 2018). The first PI(3,5)P_2_ biosensors, ML1N and a higher affinity counterpart ML1Nx2, were derived from the cation channel TRPML1 (Samie et al., 2013; Li et al., 2013). However, later studies showed that depleting PI(3,5)P_2_ did not cause the ML1Nx2 sensor to come off of endosomal membranes, indicating that off-target interactions are mediating membrane localization of the sensor (Pemberton et al., 2025; Hammond et al., 2015). Structural studies also showed that the N-terminal domain of TRPML1 used in the sensor was insufficient to bind PI(3,5)P_2_ (Hirschi et al., 2017; Chen et al., 2017). Thus, ML1Nx2 is not a useful tool to study PI(3,5)P_2_ .

A more recent biosensor, using Dictyostelium sorting nexin-like protein SnxA, which contains PI(3,5)P_2_ -binding phox homology (PX) domains, better fulfills the biosensor requirements (Vines et al., 2023). SnxA showed PI(3,5)P_2_ to be localized throughout the endosomal pathway, being somewhat enriched on Rab7 endosomes, but especially enriched on specialized endosomal membranes such as macropinosomes and phagosomes (Pemberton et al., 2025; Maib et al., 2024; Vines et al., 2023).

*In vitro*, both a biosensor made with full-length SnxA and a biosensor using the isolated PX domains in tandem (2xPx-SnxA) show specificity for PI(3,5)P_2_ : with a K _d_ of 187 nM for SnxA and 217 nM for 2xPx-SnxA. Furthermore, membrane association of either biosensor in cells depends on PI(3,5)P_2_ production (Vines et al., 2023). This work noted that within cells, 2xPx-SnxA produced more striking images with better signal to noise. Interestingly though, overexpression of the 2xPx-SnxA sensor caused some growth defects that were attributed to lipid sequestration (Vines et al., 2023). However, as there is no direct comparison of these sensors’ response to PI(3,5)P_2_ production in cells, discerning which SnxA sensor to use is not straightforward.

Later, Pemberton et al. (2025) further demonstrated using a minimal, recruitable PIKfyve construct that PI(3,5)P_2_ was sufficient to recruit SnxA to Rab5 endosomes. However, SnxA’s ability to bind to PI(3,5)P2 outside the endosomal system was not tested. As SnxA showed some basal localization at Rab5 endosomes, the possibility that SnxA binds to endosomes through coincidence detection is still unresolved. To definitively show that SnxA can unbiasedly bind to PI(3,5)P_2_ in any membrane where it may be produced, PI(3,5)P_2_ must be shown to recruit SnxA to an orthogonal membrane.

In this study, we independently validated the SnxA biosensor in live HEK293A cells. We showed the SnxA can unbiasedly report on PI(3,5)P_2_ production at membranes outside of the endosomal pathway and that it does so with more sensitivity than 2xPx-SnxA or ML1Nx2. We further confirmed that SnxA dissociates from membranes upon PI(3,5)P_2_ depletion, we used SnxA to track PI3P conversion into PI(3,5)P_2_, and we observed PI(3,5)P_2_ loss during PIKfyve inhibition. Together, these results provide independent validation of SnxA as a selective and sensitive PI(3,5)P_2_ biosensor and demonstrate its utility for tracking PI(3,5)P2 dynamics in living cells and in disease models.

## Results

### SnxA depends on PI(3,5)P2 for membrane localization

We first set out to reproduce previous results showing that SnxA depends on PI(3,5)P_2_ for its membrane localization within HEK293A cells. In other studies, this was done using deletion of PIKfyve or inhibition of PIKfyve with apilimod (Pemberton et al., 2025; Vines et al., 2023). Thus, we set out to validate this result by an independent method. We turned to the phosphatase myotubularin 1 (MTM1), which dephosphorylates both PI(3,5)P_2_ and PI3P at the 3-OH position (Schaletzky et al., 2003). We used MTM1 in a chemically inducible dimerization system using the FK506 binding protein (FKBP) and the FKBP-rapamycin-binding domain from mTOR (FRB). This approach has been used previously to deplete PI3P from Rab5 endosomes (Hammond et al., 2014, 2015).

In this assay, the FKBP-tagged MTM1 dimerizes with lysosome-targeted LAMP1-FRB after the addition of rapamycin. This results in acute depletion of PI3P and PI(3,5)P_2_ . We monitored levels of PI3P using the EEA1-FYVE biosensor (Balla et al., 2000; Burd and Emr, 1998), and quantified them by determining the fluorescence intensity of the biosensor at LAMP1-positive membranes to its intensity throughout the whole cell. This same method was used to quantify relative PI(3,5)P_2_ levels using SnxA **(Fig. 1A)**. We saw that rapamycin robustly recruited FKBP-MTM1 to the LAMP1-positive lysosomes, and the activity of MTM1 was confirmed by a substantial loss of EEA1-FYVE. We also saw the loss of SnxA at lysosomes, indicating that SnxA indeed depends on the presence of PI(3,5)P_2_ (or PI3P) for lysosome binding **(Fig. 1B)**.

**Figure 1.**
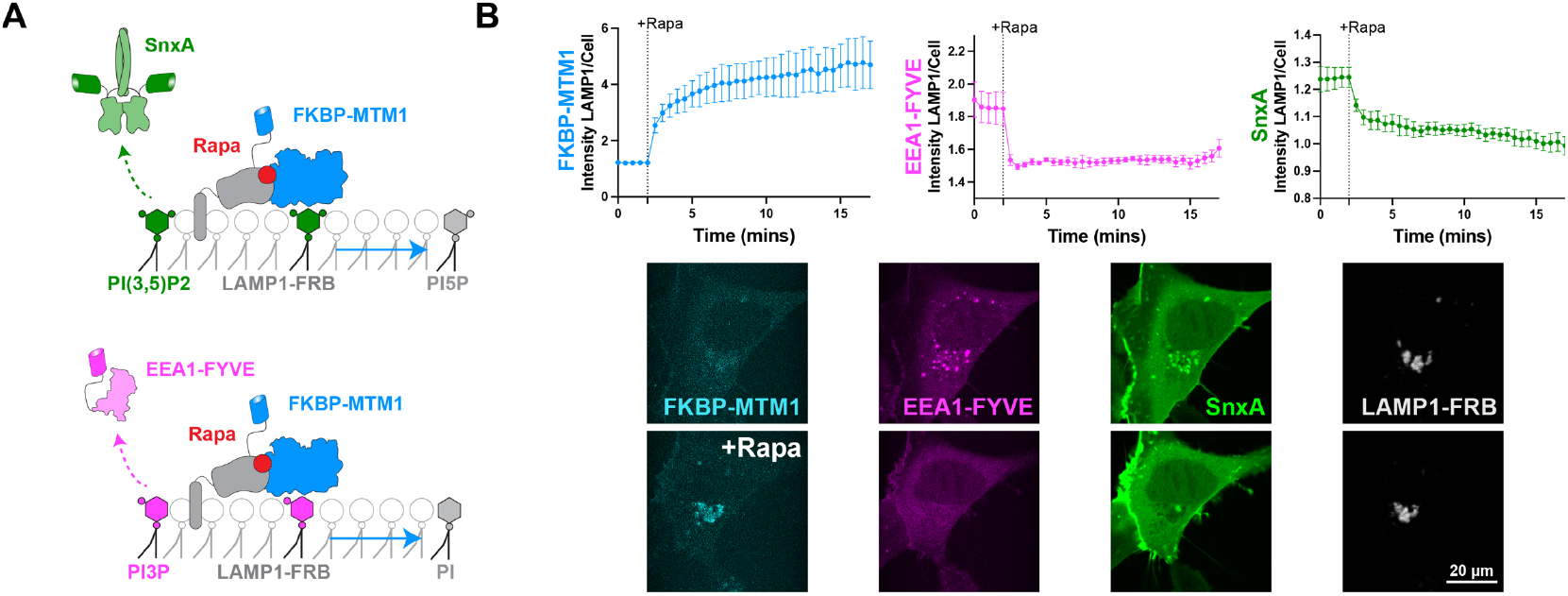
SnxA depends on PI(3.5)P2 for membrane localization. **(A)** Schematic depicting the recruitment of a chemically inducible dimerization system with MTM1 which leads to the dephosphorylation of both PI(3,5)P2 and PI3P, thereby removing SnxA and EEA1-FYVE from lysosomal membranes. **(B)** Recruitment of FKBP-MTM1 and loss of EEA1-FYVE and SnxA were quantified as the fluorescence intensity of the constructs at LAMP1-positive membranes to that within the whole cell. Data shown is the grand mean of 3 experiments ± SEM. A total of 34 cells were analyzed. Representative confocal images of the first and last time point are shown.

### PI(3,5)P2 loss precedes vacuolation during apilimod treatment

As MTM1 depletes two lipid species that SnxA could possibly be interacting with, we next turned to apilimod treatment to inhibit PIKfyve and PI(3,5)P_2_ production specifically. We stained lysosomes/late endosomes (LyLE) with fluorescent dextran loading and then took z-stacks of the cells to image a large volume of endosomes during apilimod treatment. After about 4-6 minutes of apilimod treatment, nearly all of the SnxA was removed from lysosomes/late endosomes **(Fig. 2A)**, in agreement with previous data (Pemberton et al., 2025; Vines et al., 2023). Then after around 30 minutes, we saw significant vacuolation, indicating that PIKfyve was inhibited as expected **(Fig. 2B)**. This data supports **Fig. 1** in showing that SnxA depends on PI(3,5)P_2_ for membrane localization.

**Figure 2.**
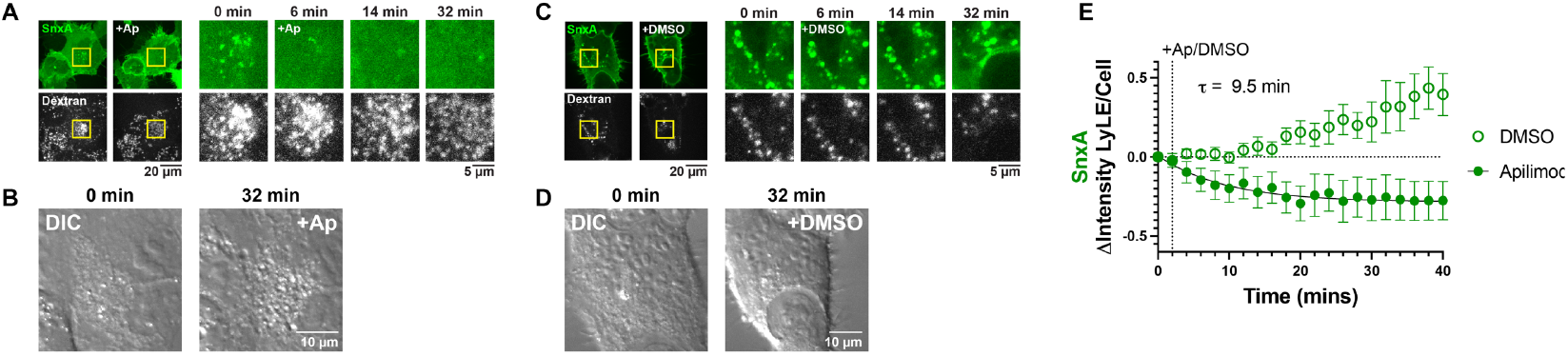
PI(3,5)P2 loss precedes vacuolation during apilimod treatment. **(A)** Representative mean intensity projections of 11 0.175 µm equatorial confocal sections showing the loss of SnxA’s vesicular localization a few minutes after treatment with 1 µM apilimod. **(B)** Differential interference contrast images (DIC) show that vacuolation does not occur until 25-30 minutes after apilimod treatment. **(C)** Representative confocal projections as in A showing that treatment with 0.02% DMSO does not change SnxA localization. **(D)** DIC images showing that 0.02% DMSO does not cause vacuolation. **(E)** Quantification of SnxA localization at lysosomes/late endosomes (LyLE) after treatment with 1 µM apilimod or 0.02% DMSO. A one phase decay nonlinear fit was calculated for the apilimod treatment. K was constrained to > 0, which produced a time constant (t) of 9.507 min with a 95% confidence interval of 7.043 to 13.03 min. (DF = 29, R2 = 0.8856). Graph shows the mean ± SEM for 14-16 cells from 3 independent experiments.

In the DMSO control dish, SnxA remained stably associated with the lysosomes/late endosomes, and no vacuolation occurred **(Fig. 2C,D)**. When the loss of SnxA with apilimod was quantified, fitting a one phase decay model determined that SnxA came off of the membranes with a time constant (τ) of 9.5 minutes, which is well before vacuolation begins **(Fig. 2E)**.

The quick depletion of PI(3,5)P_2_ that we measured by SnxA after apilimod treatment is in agreement with data obtained using the PIKfyve inhibitor YM201636. Vacuolation was visualized 35 minutes after the addition of the inhibitor, but HPLC data showed that PI(3,5)P_2_ levels were depleted to their minimum level in only 2.5 minutes (Jefferies et al., 2008; Zolov et al., 2012). This experiment further demonstrates the effectiveness of SnxA as this biosensor allowed us to confirm biochemical kinetic data in real-time and with live cells.

### PI(3,5)P_2_ does not significantly overlap with PI3P

When imaging cells, we noticed that often SnxA and the PI3P biosensor EEA1-FYVE do not overlap. To further define this relationship between the endosomal lipids, we analyzed the images taken throughout this study, before any manipulations were done, to see if there was co-localization between SnxA and EEA1-FYVE. While we see some SnxA and EEA1-FYVE in close proximity to each other, we don’t often see co-localization of the two. Furthermore, we can find several discrete examples of vesicles only marked with SnxA, as shown by white arrows **(Fig. 3A)**.

**Figure 3.**
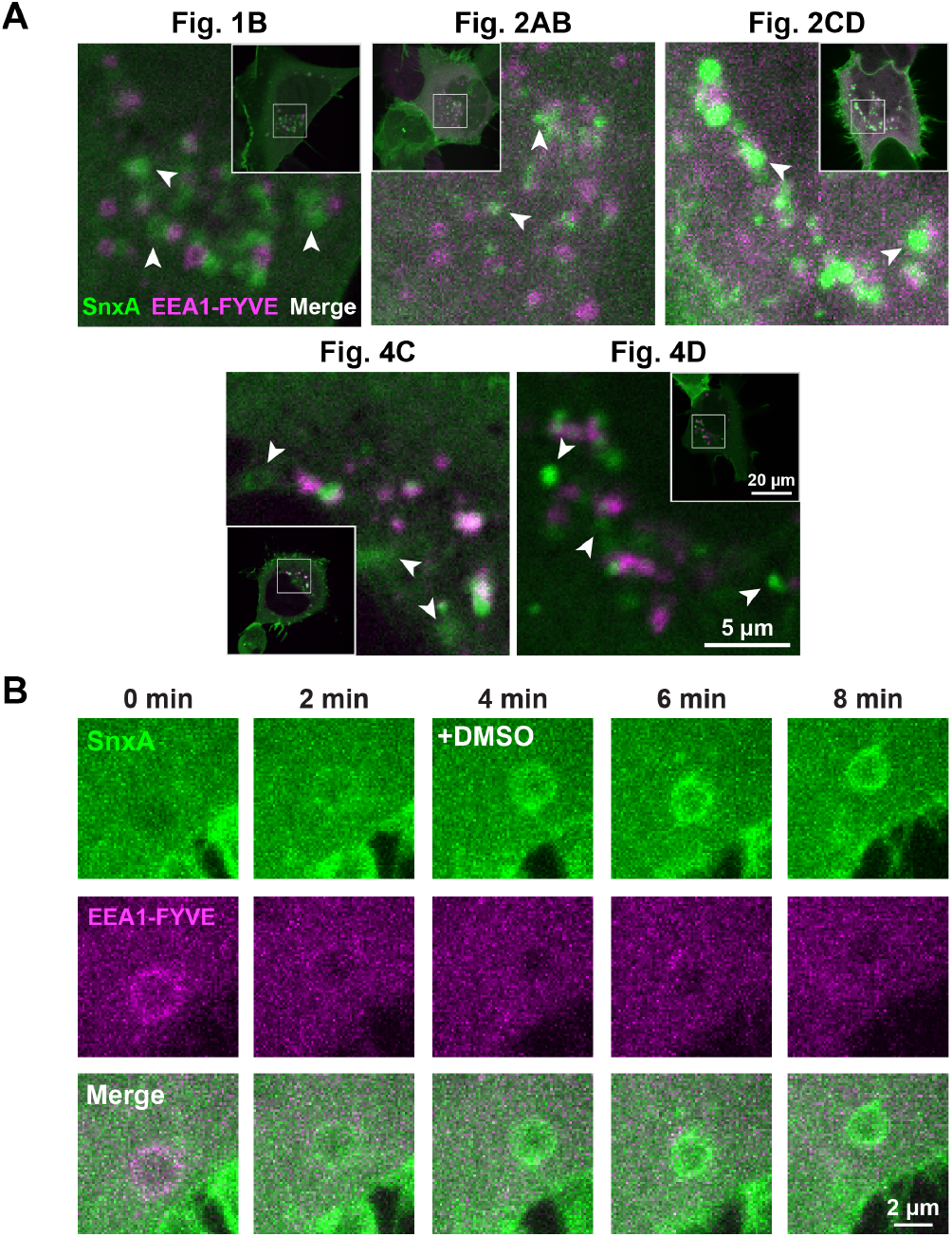
PI(3,5)P_2_ does not significantly overlap with PI3P. **(A)** Confocal images of cells used throughout this study expressing both SnxA and EEA1-FYVE. The white arrows show examples of where the biosensors don’t co-localize. The figure where each cell is duplicated from is as noted. **(B)** Timelapse images of a representative vesicle showing the transition from EEA1-FYVE labelling to SnxA labelling in an untreated cell expressing both biosensors.

Additionally, in the z-stack images, we saw several good examples of vesicles that switched from being EEA1-FYVE-labelled to being SnxA-labelled. First, this indicated the conversion of PI3P to PI(3,5)P_2_ and thus increased our confidence that SnxA is responding to PI(3,5)P_2_ . Second, this transition was fairly quick, happening in about 4 minutes. Notably, it only occurred in a small subset of EEA1-FYVE-positive vesicles **(Fig. 3B)**. This quick transition could account for the lack of SnxA and EEA1-FYVE overlap seen in static images. Overall, this data demonstrates that the SnxA sensor can provide important insights into the relationship between PI(3,5)P_2_ and the other phosphoinositides.

### Design of a recruitable PIKfyve construct

In the recent study characterizing SnxA, Pemberton et al. designed a minimal catalytic fragment of PIKfyve that could be used in the FKBP-FRB system (Pemberton et al., 2025). While that manuscript was in preparation, we had independently designed the same construct, using AlphaFold3 (Abramson et al., 2024) with some trial and error to determine which domains of the full-length murine PIKfyve were necessary to retain activity. We found that the conserved cysteine-rich (CCR) domain was packed against the kinase domain and therefore likely impacted folding or stability of the kinase domain. However, the CCR itself could only properly fold when the 3-helix bundle was present. Then, for use in our imaging experiments, we also included FKBP and a BFP fluorophore with some flexible linkers. Using AlphaFold3, we saw that the structures of the PIKfyve domains in our construct resembled the folded domain structures of full-length PIKfyve. As this construct contains the kinase domain and the CCR, we named this construct FKBP-PIKfyve-CCR-Kina **(Fig 4A)**. We then added mutations (K1876E and V1883I in full-length *M. musculus* PIKfyve) that inactivate PIKfyve to create FKBP-PIKfyve-CCR-Kina-Dead as a control.

**Figure 4.**
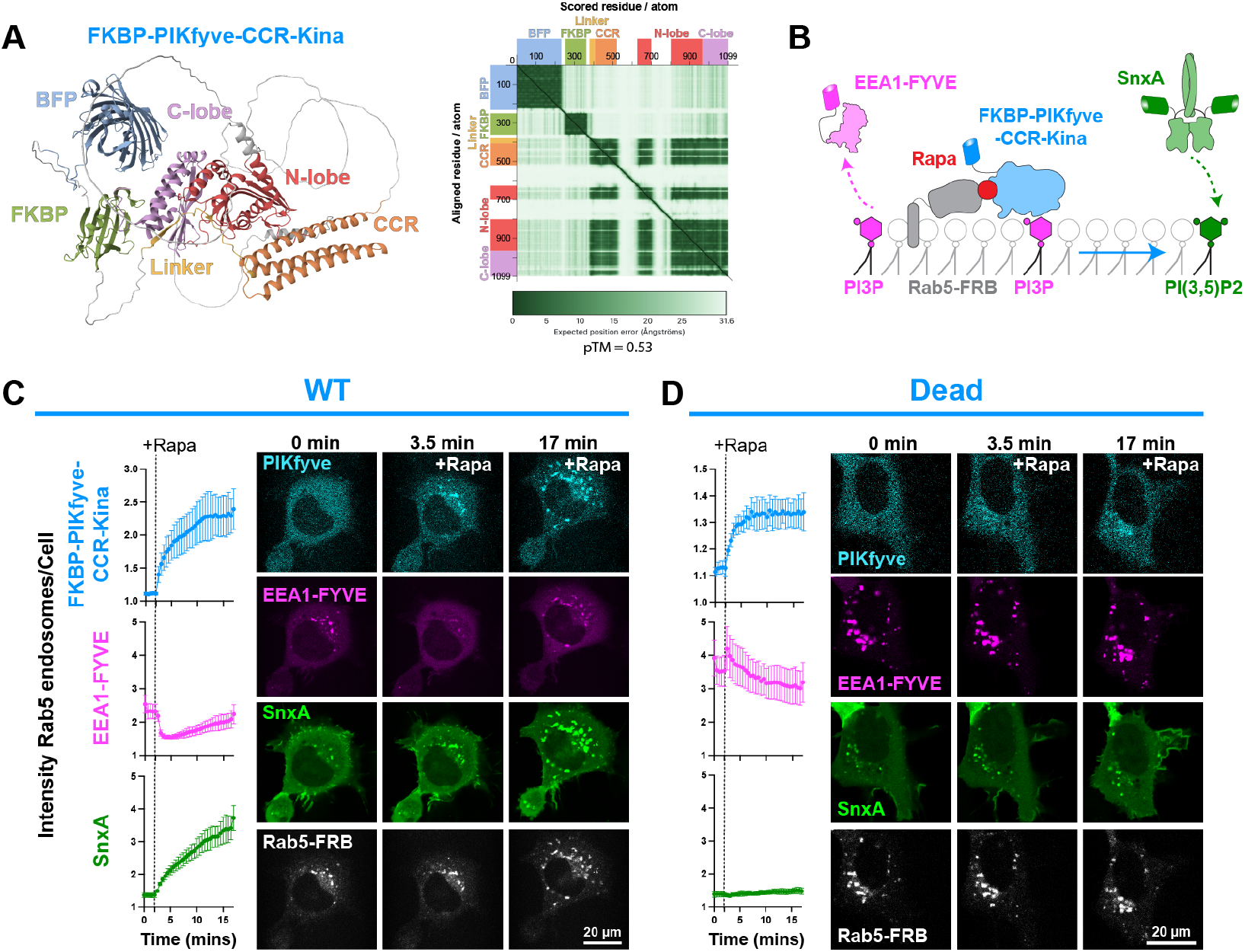
Design of a recruitable PIKfyve construct. **(A)** The AlphaFold3 predicted structure of the FKBP-PIKfyve-CCR-Kina construct. Both the conserved Cys-rich region (CCR) and the kinase domain, made up of the N-lobe, C-lobe, and linker, were included to facilitate proper domain folding. The predicted aligned error (PAE) is shown to demonstrate high confidence in the predicted structures of the defined domains. The predicted template modeling (pTM) score is given. Scores above 0.5 indicate confidence that the predicted structure aligns with the true structure. **(B)** Schematic showing how FKBP-PIKfyve-CCR-Kina activity was monitored by EEA1-FYVE and SnxA biosensors after the PIKfyve construct was recruited to Rab5 endosomes with rapamycin. **(C)** Recruitment of the WT PIKfyve-CCR-Kina to Rab5 endosomes and changes in biosensor localization at these membranes is shown by the fluorescence intensity ratios of the constructs at Rab5 endosomes over the whole cell. Data shown is the grand mean ± SEM of 3 experiments. A total of 35 cells were analyzed. Confocal images over time are shown to the right. **(D)** As in C, but now utilizing the catalytically dead K1876E/V1883I mutant of the FKBP-PIKfyve-CCR-Kina as a negative control. Data shown is the grand mean ± SEM for 3 experiments. A total of 39 cells were analyzed.

We first used the FKBP-PIKfyve-CCR-Kina to produce PI(3,5)P_2_ on Rab5 endosomes, where its substrate PI3P endogenously resides **(Fig. 4B)**. Rapamycin recruited FKBP-PIKfyve-CCR-Kina to Rab5-positive membranes, and we saw a subsequent depletion of PI3P shown by loss of EEA1-FYVE. The resulting PI(3,5)P_2_ production then proved to be sufficient for SnxA recruitment, as we saw very robust labeling of Rab5 endosomes with SnxA **(Fig. 4C)**. When FKBP-PIKfyve-CCR-Kina-Dead was expressed as a control, we did not see a significant change in either EEA1-FYVE or SnxA localization, indicating that the changes in the biosensors seen previously were dependent on PIKfyve activity **(Fig. 4D)**.

### Production of PI(3,5)P_2_ at orthogonal membranes is sufficient to recruit SnxA

Next, to fully confirm that PI(3,5)P_2_ is sufficient for SnxA recruitment, we targeted the FKBP-PIKfyve-CCR-Kina constructs to the mitochondrial membrane, which lacks PI3P and so presumably, also lacks typical endosomal effector proteins. Therefore, producing PI(3,5)P_2_ at this membrane is a more rigorous way to show that PI(3,5)P_2_ is sufficient for SnxA membrane association. To make PI(3,5)P_2_ at mitochondria, we co-expressed FKBP-MavQ along with the FKBP-PIKfyve-CCR-Kina. MavQ is a *L. pneumophila* effector that produces PI3P on the ER through the phosphorylation of phosphatidylinositol (PI) (Hsieh et al., 2021). Therefore, we recruited FKBP-MavQ to mitochondrial membranes, so it could supply the FKBP-PIKfyve-CCR-Kina with its PI3P substrate **(Fig. 5A)**.

**Figure 5.**
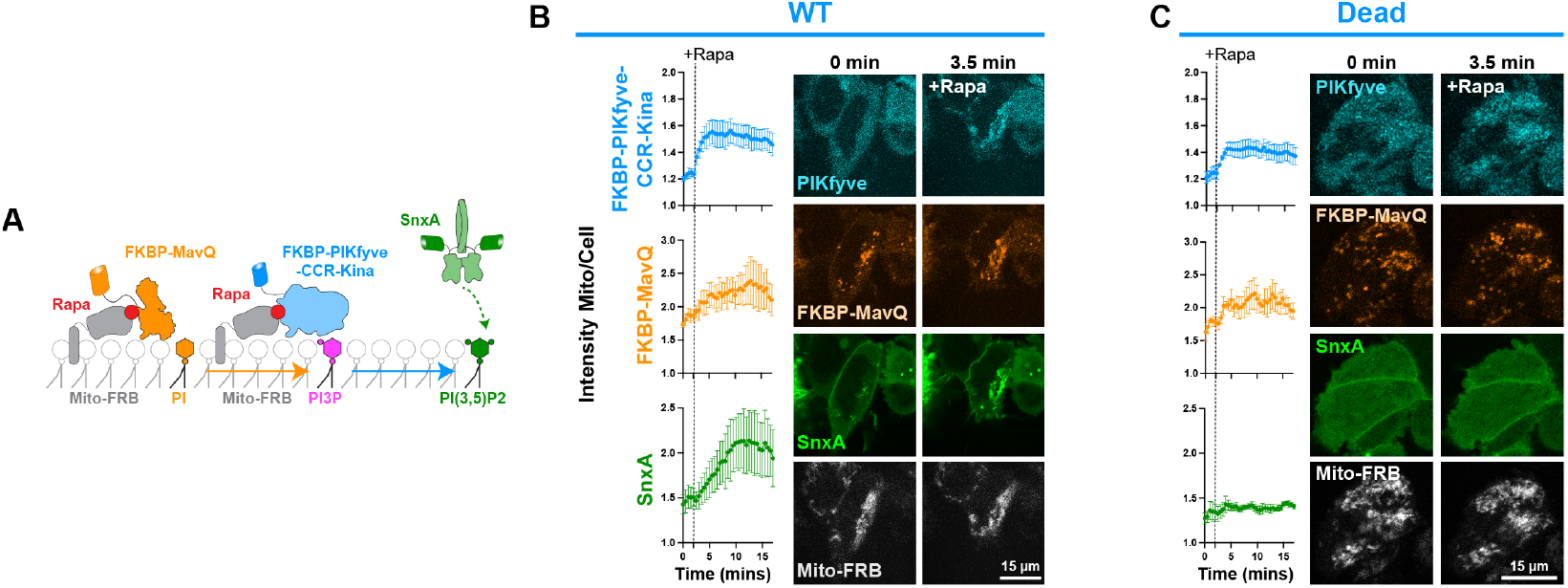
Production of PI(3,5)P_2_ at orthogonal membranes is sufficient to recruit SnxA. **(A)** Schematic showing ectopic PI(3,5)P_2_ production on mitochondrial membranes through co-expression and co-recruitment of FKBP-MavQ and FKBP-PIKfyve-CCR-Kina. **(B)** Quantification of the WT PIKfyve, MavQ, and SnxA at the mitochondria using the ratio of fluorescence at the mitochondria to that within the whole cell. Data shown is the grand mean ± SEM of 3 experiments. A total of 33 cells were analyzed. Representative confocal images of the time course are shown to the right. **(C)** As in B, but with the kinase dead mutant of the FKBP-PIKfyve-CCR-Kina. Data shown is the grand mean of 3 experiments ± SEM. A total of 29 cells were analyzed.

Interestingly, FKBP-MavQ already showed some co-localization with mitochondrial membranes even before rapamycin addition **(Fig. 5B,C)**. However, we did see rapamycin induce recruitment of FKBP-PIKfyve-CCR-Kina and increase recruitment of FKBP-MavQ. The resulting production of PI(3,5)P_2_ then caused robust localization of SnxA at the mitochondria **(Fig. 5B)**, which was not seen when the FKBP-PIKfyve-CCR-Kina-Dead was used **(Fig. 5C)**. Overall, this data increased our confidence that SnxA is a robust PI(3,5)P_2_ biosensor.

### SnxA outcompetes 2xPx-SnxA and ML1Nx2 for PI(3,5)P_2_ binding

Now having seen that SnxA fully meets the criteria for a PI(3,5)P_2_ biosensor, we wanted to show its sensitivity as compared to the other PI(3,5)P_2_ biosensors 2xPx-SnxA and ML1Nx2, as direct comparison of the responses of these sensors to PI(3,5)P_2_ has not been done previously.

We first co-expressed SnxA and 2xPx-SnxA in the same cell to simultaneously monitor their response to PI(3,5)P_2_ production at Rab5 endosomes. 2xPx-SnxA showed strong nuclear localization as described previously. Though this nuclear accumulation can be circumvented through the addition of a nuclear export sequence (Pemberton et al., 2025). After recruitment of FKBP-PIKyve-CCR-Kina, the 2xPx-SnxA sensor labelled the Rab5 endosomes. In comparison, SnxA showed some slight enrichment at Rab5 endosomes under basal conditions, and then more robustly translocated to Rab5 endosomes after rapamycin addition **(Fig. 6A)**. Importantly, neither biosensor was enriched at Rab5 endosomes when FKBP-PIKfyve-CCR-Kina-Dead was recruited there **(Fig. 6B)**. Altogether, this data aligns with the reported high affinity for SnxA and suggests that SnxA is outcompeting 2xPx-SnxA for PI(3,5)P_2_ binding.

**Figure 6.**
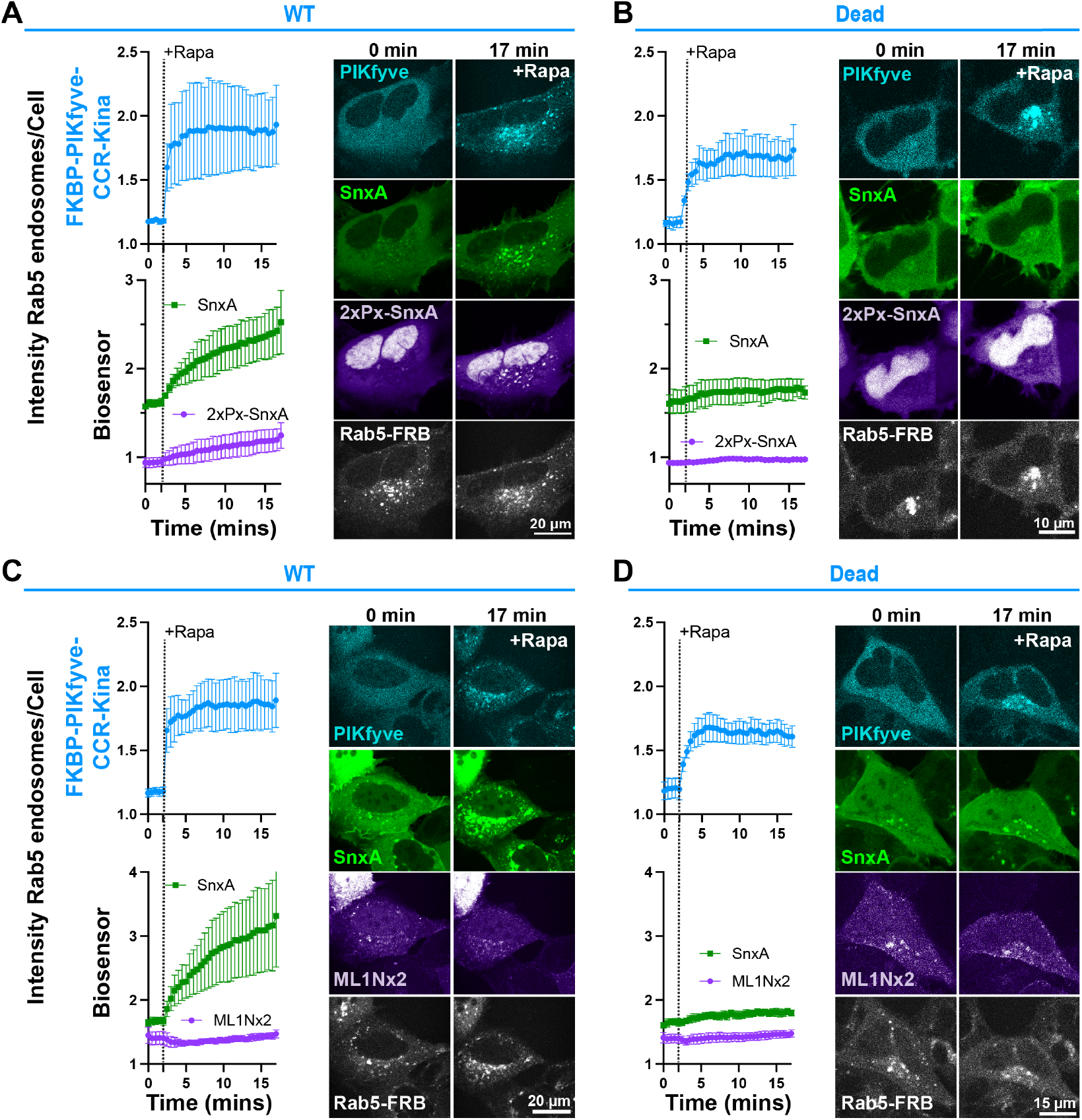
Comparison of SnxA, 2xPx-SnxA and ML1Nx2 for ectopic PI(3,5)P_2_ binding. **(A)** Recruitment of the FKBP-PIKfyve-CCR-Kina to Rab5 membranes and the subsequent translocation of SnxA and 2xPx-SnxA biosensors was quantified as a fluorescence intensity ratio of fluorescence at Rab5 endosomes to that within the whole cell. Data shown is the grand mean ± SEM of 3 independent experiments. A total of 34 cells were analyzed. Representative confocal images of the first and last time point are shown to the right. **(B)** The kinase dead mutant of the FKBP-PIKfyve-CCR-Kina construct was used in an experiment parallel to that in A as a negative control. Data shown is the grand mean ± SEM of 3 independent experiments. A total of 38 cells were analyzed. **(C)** The SnxA biosensor was co-expressed with the ML1Nx2 biosensor to compare their responses to PI(3,5)P_2_ production at Rab5 endosomes by FKBP-PIKfyve-CCR-Kina. Data shown is the grand mean ± SEM of 3 independent experiments. A total of 36 cells were analyzed. **(D)** The control experiment for C, showing the response of SnxA and ML1Nx2 to the kinase dead FKBP-PIKfyve-CCR-Kina at Rab5 endosomes. Data shown is the grand mean ± SEM of 3 independent experiments. A total of 36 cells were analyzed.

We next tested ML1Nx2 against SnxA by co-expressing them in the same cells. In this case, we did not see ML1Nx2 respond to PI(3,5)P_2_ production by FKBP-PIKfyve-CCR-Kina at Rab5 endosomes, while SnxA showed very strong recruitment **(Fig. 6C)**. The SnxA translocation was lost when the catalytically dead PIKfyve construct was used **(Fig. 6D)**. This data agrees with past evidence that suggested that ML1Nx2 targets endosomes through off-target interactions, and therefore PI(3,5)P_2_ is not sufficient to recruit ML1Nx2 to endosomal membranes (Hammond et al., 2015). Although, when the ML1Nx2 sensor was converted into a more sensitive BRET-based probe, a very small response to PI(3,5)P_2_ production on Rab5 endosomes was seen (Pemberton et al., 2025). Overall though, our data and previous results all indicate that SnxA and 2xPx-SnxA have far superior sensitivity and selectivity in comparison to ML1Nx2. Thereby, use of the SnxA-based biosensors over ML1Nx2 is highly recommended.

## Discussion

In this study, we showed that SnxA membrane localization depends on PI(3,5)P_2_ **(Fig. 1, 2)** and that PI(3,5)P_2_ production by a minimal PIKfyve construct, PIKfyve-CCR-Kina, is sufficient to recruit SnxA to mitochondrial membranes, which aren’t expected to contain other endosomal lipids or effectors **(Fig. 4, 5)**. Finally, we demonstrated that SnxA responds to PI(3,5)P_2_ production more robustly than the 2xPx-SnxA and ML1Nx2 sensors **(Fig. 6)**. Overall, our work validated SnxA as a selective and sensitive PI(3,5)P_2_ biosensor. We showed that SnxA is suitable for live-cell studies of PI(3,5)P_2_ dynamics, as SnxA allowed us to visualize loss of PI(3,5)P_2_ during apilimod treatment and its conversion from PI3P **(Fig. 2, 3)**. SnxA will thus be instrumental in future studies investigating PI(3,5)P_2_’s regulation in normal cellular physiology and in disease contexts.

Importantly, our study is the first to show that SnxA responds to PI(3,5)P_2_ produced in an orthogonal membrane outside of endosomal pathway **(Fig. 5)**. While previous studies showed that SnxA localization at Rab5 endosomes was decreased with PI(3,5)P_2_ depletion and increased with PI(3,5)P_2_ synthesis, these experiments did not address whether SnxA was binding to Rab5 endosomes through coincidence detection of PI(3,5)P_2_ and another lipid or protein (Vines et al., 2023; Pemberton et al., 2025). However, by showing SnxA recruitment to mitochondria upon PI(3,5)P_2_ production, we have definitively shown that PI(3,5)P_2_ is sufficient for SnxA to bind to membranes.

The ability of a biosensor to unbiasedly report on lipid levels is a key feature of a reliable biosensor, as it ensures that the biosensor will detect the lipid in any membrane that the lipid may be produced in, rather than only “seeing” a specific pool of lipid. For example, PI4P was detected only in the Golgi by PH-FAAP1 biosensors, which localize to the Golgi by interactions with both PI4P and Arf1. However, the P4M biosensor, which was shown to detect ectopic PI4P made in the ER, and therefore bind to PI4P but not Arf1, also revealed PI4P pools in endosomes, lysosomes, and at the PM (Hammond et al., 2014).

Furthermore, this work directly compared SnxA, 2xPx-SnxA, and ML1Nx2 by expressing the sensors in the same cell **(Fig. 6)**. Pemberton et al. compared 2xPx-SnxA to the monomeric probe Px-SnxA, but not the full-length SnxA probe (Pemberton et al., 2025). Vines et al. showed SnxA to have higher affinity for PI(3,5)P_2_ *in vitro* compared to 2xPx-SnxA, and SnxA showed more basal vesicular localization compared to 2xPx-SnxA, but this study only used static imaging (Vines et al., 2023). We therefore directly compared 2xPx-SnxA to SnxA over time within the same cell. Our data agreed with SnxA having higher affinity for PI(3,5)P_2_, as we saw SnxA have more basal localization at Rab5 endosomes and a more robust response to PI(3,5)P_2_ . Additionally, we confirmed previous data showing that ML1Nx2 does not accurately respond to PI(3,5)P_2_ levels.

While this study confirmed SnxA’s high sensitivity, our data does not address if SnxA sequesters PI(3,5)P_2_ and disrupts its interactions with effector proteins. This consideration is especially important as PI(3,5)P_2_ is a very low abundance lipid. It is estimated to be about 0.038% of the total phosphoinositides in mouse embryonic fibroblasts (Steinfeld et al., 2021). At these low levels, it is a real concern that the overexpressed biosensor will bind to all of the available lipid and sequester it from effector proteins. Aligning with this idea, expression of the 2xPx-SnxA biosensor has been previously associated with growth defects, which are potentially a result of disrupted lipid signaling (Vines et al., 2023). On the other hand, it is somewhat reassuring that extensive vacuolation of the cell is not seen with SnxA or 2xPx-SnxA expression; if the probes completely sequestered PI(3,5)P_2_, we would expect the same phenotype as PIKfyve inhibition.

One way to circumvent lipid sequestration is to use a lower affinity biosensor that will dissociate from the lipid more easily. However, a lower affinity biosensor using a single PX domain (Px-SnxA) did not show *in vitro* binding to PI(3,5)P_2_ or any basal endosomal localization (Pemberton et al., 2025; Vines et al., 2023). Therefore, we suggest that expressing the full-length SnxA biosensor at lower levels could minimize competition with effector proteins. We have seen this approach work with biosensors for PIP_3_ (Holmes et al., 2025).

One of the most important uses of SnxA will be in the study of PIKfyve-related diseases. PIKfyve dysfunction has been linked to several diseases, yet the role of PI(3,5)P_2_ in disease pathology is still unclear. PIKfyve-generated PI(3,5)P_2_ is the precursor for phosphatidylinositol 5-phosphate (PI5P) by way of MTM-mediated dephosphorylation of PI(3,5)P_2_ (Zolov et al., 2012). Therefore, the effects of PI(3,5)P_2_ loss are intertwined with the effects of PI5P loss in PIKfyve-related diseases. Notably, there is no characterized biosensor for PI5P, so before SnxA’s discovery, distinguishing between the role of PI(3,5)P_2_ and PI5P in cellular events has been difficult. Furthermore, the relationship between PIKfyve and levels of these lipids is not straightforward in disease states. Mouse embryonic fibroblasts with only one PIKfyve allele show a 60% reduction in PI(3,5)P_2_ and PI5P levels despite only a 50% decrease in PIKfyve protein levels (Ikonomov et al., 2011). Therefore, the fully validated SnxA biosensor will be invaluable for detailed studies of PI(3,5)P_2_ in disease models.

In this work, SnxA has already corroborated important information for our understanding of PIKfyve-related diseases. SnxA confirmed that PI(3,5)P_2_ depletion precedes cellular changes. Apilimod treatment eliminated SnxA’s association with late endosomes and lysosomes within 9.5 minutes, whereas endosomal vacuolation began only after 25– 30 minutes **(Fig. 2)**. The time course of PIKfyve inhibition gradually leading to vacuolation is well-documented (Choy et al., 2018; Sbrissa et al., 2018). Additionally, a rapid loss of PI(3,5)P_2_ with PIKfyve inhibition ( < 5 mins) has been determined biochemically (Zolov et al., 2012). However, when the ML1Nx2 biosensor was used to monitor PI(3,5)P_2_ loss with the inhibitor YM201636, a gradual decrease in PI(3,5)P_2_ was seen. Significant loss of PI(3,5)P_2_ occurred after 3 hours of inhibitor treatment and further loss was seen with overnight treatment (Li et al., 2013). Here we show that the SnxA sensor, which better fulfills the biosensor criteria as compared to ML1Nx2, is more in agreement with the biochemical data than the ML1Nx2 studies were. This shows the power of SnxA; it allowed us to confirm PI(3,5)P_2_ kinetics in real-time that had previously only been seen biochemically.

Additionally, the lagging appearance of large vacuoles after PI(3,5)P_2_ reduction suggests that the lipid regulates endosome size through a delayed mechanism, but the exact mechanism underlying this phenotype is still unclear. One model hypothesizes that PIKfyve inhibition leads to vesicle swelling through changes in ion homeostasis and osmosis. For example, PI(3,5)P_2_ activation of two pore channels (TPCs) causes Na ^+^ efflux from the lysosome (Wang et al., 2012). In macropinosomes, this TPC-dependent ion movement results in loss of water and vesicle shrinkage (Freeman et al., 2020). Therefore, the increase in vacuolar size with PIKfyve inhibition could be due to the inability of the TPCs to drive water efflux from the vesicles. In support of this hypothesis, applying a hyperosmotic solution to PIKfyve-inhibited cells reversed the swelling of the vacuole (Freeman et al., 2020).

Similarly, PI(3,5)P_2_ was shown to regulate Cl^-^/H ^+^ exchangers in endosomes. The voltage-gated chloride channel 3 (ClC-3) regulates vacuolation of endosomes by increasing luminal Cl ^-^ levels, which causes osmotic swelling (Sonawane et al., 2002). However, ClC-3 can be inhibited by a regulatory T9A subunit blocking the ion pore. PI(3,5)P_2_ was shown to stabilize the association of ClC-3 and T9A. Apilimod-induced depletion of PI(3,5)P_2_ overrode T9A overexpression to cause constitutive ion influx and ClC-3-mediated vacuolation (Schrecker et al., 2025).

In another model, the loss of PI(3,5)P_2_ affects endosomal pH. This occurs through dysregulation of v-ATPases causing improper endosomal/vacuole acidification (Banerjee and Kane, 2020; Gary et al., 1998; Yamamoto et al., 1995). While it is still unclear how pH regulates vacuolation under PIKfyve inhibition, it is thought that proper pH levels are needed to maintain endosomal identity and regulate fission or fusion events or that a tightly controlled pH is needed for proper ion homeostasis and water flux. Therefore, upon PIKfyve inhibition, enlarged endosomes are speculated to occur due to dysregulated pH levels leading to increased fusion or increased water influx. Changing the pH of endosomal compartments has been seen to reverse vacuolation (Nicoli et al., 2019), but other studies show that pH does not affect vacuole size (Leray et al., 2022). Therefore, more work needs to be done to determine the exact relationship between pH and endosomal size and to uncover how PIKfyve inhibition alters it.

Finally, PIKfyve inhibition is thought to increase the size of endosomes through membrane remodeling mechanisms. In these cases, inhibiting PIKfyve activity disrupts the balance of lysosomal fusion and fission events (Choy et al., 2018), or reduces the capacity of endolysosomes to form lysosomes (Bissig et al., 2017). Both of these mechanisms result in the stalling of endosomal trafficking where endosomes gain volume from upstream trafficking events but then don’t dissipate that volume through further trafficking or fission events.

While any of these mechanisms could explain the delay in the appearance of the endosomal phenotype after rapid PI(3,5)P_2_ loss, all of them will need further testing. Our ability to test these models is now more achievable using the SnxA biosensor and the FKBP-FRB systems that we describe here. Using these tools in combination with live-cell imaging will allow us to determine the effects of acute changes in PI(3,5)P_2_ on things like endosomal ion levels or fusion events.

The SnxA biosensor will also help us uncover more detail about PI(3,5)P_2_ in the context of the other phospholipids. There was evidence that PI(3,5)P_2_ was synthesized from PI3P since the first discovery of PI(3,5)P_2_ (Dove et al., 1997) . However, there has been some conflicting data as to the localization of PI(3,5)P_2_ relative to that of PI3P. Some early studies suggested that PIKfyve and PI3P localize in separate membrane domains within early endosomes (Cabezas et al., 2006), which brings up the question: how much overlap is there between PI3P and PI(3,5)P_2_ in the endosomal pathway? Consistent with prior work (Vines et al., 2023; Pemberton et al., 2025), we observed little overlap between the two lipids in static images of a population of endosomes **(Fig. 3)**. But, we could observe a short temporal overlap between the two lipids within a single endosome, lasting only a few minutes. Furthermore, we observed that this transition happens spontaneously in cells **(Fig. 3)**, and we did not see any evidence of lipid-specific membrane domains. This short window of co-localization will be important to consider as we continue to elucidate the functions of PI(3,5)P_2_ effector proteins, some of which like WIPI49 bind to both PI3P and PI(3,5)P_2_ (Jeffries et al., 2004).

Overall, we have found SnxA to be a selective and sensitive biosensor that fills a critical gap in the tools available to study the sparse but essential lipid PI(3,5)P_2_.

## Author Contributions

Data analysis; Investigation: Visualization; Writing – Review & Editing: Tiernan Swayhoover. Conceptualization; Visualization; Writing - Original Draft; Writing - Review & Editing: Claire C. Weckerly. Conceptualization; Data analysis; Writing - Review & Editing: Gerald R.V. Hammond. The authors declare no competing financial interests.

## Materials and Methods

### Cell Culture and Transfection

HEK293A cells (Invitrogen R70507) were maintained at 37º C and 5% CO _2_ . They were passaged 1:5 twice weekly using the following protocol: flasks were washed with 10 mL 1x PBS (Fisher Scientific BP3994), 1 mL TrypLE (Life Technologies 12604039) was used to dissociate the cells, and 4 mL complete Dulbecco’s modified Eagle’s media (DMEM; Life Technologies 10567022, supplemented with 10% HI-FBS (Life Technologies A5669801), 1% 100 u/mL penicillin and 100 µg/mL streptomycin (Life Technologies 15140122), 0.1% chemically defined lipid supplement (Life Technologies 11905031)) was added to the flask and the cells were resuspended by pipetting. The suspended cells were then seeded into 35 mm glass bottom dishes with a 20 mm aperture (Fisher Scientific D35-20-1.5-N). Dishes were pre-coated with 10 µL 1 mg/mL ECL cell attachment matrix (Sigma 08-110) diluted in 0.5 mL incomplete DMEM. The coating was incubated on the dishes for 30+ minutes at 37º C. After the incubation period, the coating solution was replaced with 2 mL pre-warmed complete DMEM. The cells were added into this media, seeding enough cells that the dishes would be 100% confluent on the day of imaging. The cells were then left to settle onto the dish for 2+ hours at 37º C before transfecting.

To transfect each dish of cells, 1 µg of DNA (the needed plasmids were mixed to obtain 1 µg total) was diluted in 100 µL of Opti-MEM (Life Technologies 51985091). Then, 3 µL Lipofectamine2000 (Life Technologies 11668019) was mixed in 100 µL Opti-MEM. These two solutions were mixed together and then incubated for 5-20 minutes at room temperature. 200 µL of the resulting solution was added to each dish of cells. The dishes were swirled to mix and then incubated at 37º C for 4 hours. At this time, the transfection solution was removed and replaced with 2 mL Complete Hepes-buffered Imaging Media (CHIM; 10% HI-FBS, 1% Glutamax (Life Technologies 35050061), 0.1% chemically defined lipid supplement, 2.5% 1 M Na-HEPES buffer at pH 7.4 (VWR EM-5320) diluted in 500 mL Fluorobrite media (Life Technologies A1896702)). The cells were then kept in the incubator overnight and imaged the next day.

### Dextran Labeling

For cells labelled with dextran, the transfection mixture was replaced with 2 mL complete DMEM rather than CHIM the night before. Then the day of imaging, a solution of 100 µg/mL Alexa Fluor™ 647 Dextran (ThermoFisher D22914) was made up in CHIM. Note that the dextran solution must be protected from light as much as possible. The media on the cells was aspirated, and 500 µL of the dextran solution was added to the glass inset of each dish. The cells were incubated in the dextran for 1 hour at 37º C. After the incubation, the cells were washed with 1 mL CHIM, the wash was aspirated, and then 2 mL CHIM was added to the dish for imaging.

### Plasmids and Cloning

The plasmids used in this study are listed in **Table 2**. Plasmids were cloned using PCR and the NEBuilder HiFi DNA Assembly Cloning Kit (New England Biolabs E5520S) or restriction digest (FastDigest Thermo Scientific) followed by ligation (T4 DNA Ligase, Thermo Scientific EL0011). Vector and insert sequences were either ordered as a custom gene block (IDT) or isolated using primers for an existing plasmid. Proper plasmid synthesis was confirmed by Sanger sequencing over the insertion sites and/or by long-read nanopore sequencing over the whole plasmid (GENEWIZ from Azenta Life Sciences).

### Confocal Imaging

Imaging was done using a Nikon A1R confocal microscope with a Nikon TiE inverted stage and a heated stage incubator to keep the dishes at 37º C (Tokai Hit). A dual fiber-coupled laser LU-NV launch was used to excite the cells: 405 nm for blue fluorophores, 488 nm for green fluorophores, 561 nm for red fluorophores, and 640 nm for farred fluorophores. Channel series was used so that cells were excited with 405/561 nm lasers in one pass, and 488/640 nm lasers in a separate pass in order to avoid crosstalk. Emission was collected by 500-550 nm (green), 663-737 nm (far-red), 570-620 nm (red), and 425-475 nm (blue) fibers. A 100x 1.45 NA plan apochromatic oil immersion objective was used, and the pinhole opening was set to 1.2x the Airy disc size of the longest wavelength being imaged.

To take the images, 8x line averaging, a 512-pixel scan area size, and 2x zoom was used. In some experiments, the Nikon Elements denoising software was also used. To image, first 10+ cells appropriately expressing the plasmids were found and their positions marked. A timelapse was then set to take images every 30 seconds for the total duration as noted in the figure legends. After 2 minutes of baseline imaging, the cells were stimulated with 500 µL of the solutions described in **Table 1**. Solutions were made up at a concentration 5x of that listed in **Table 1** since adding 500 µL solution to the 2 mL of CHIM on the cells resulted in a 1:5 dilution.

**Table 1.** Cell stimulations used throughout this study.

| Reagent | Manufacturer | Catalog Number | Stock solution | Concentration used on cells (diluted in CHIM) | Figures Used |
| --- | --- | --- | --- | --- | --- |
| Rapamycin | EMD | 80054-244 | 1 mM | 1 μM | Fig. 1, 4, 5, 6 |
| Apilimod | Sigma-Aldrich | SML2974-5MG | 5 mM | 1 μM | Fig. 2 |
| DMSO | Sigma | D2653-5X5ML | 100% | 0.02% | Fig. 2, 3 |

**Table 2.**
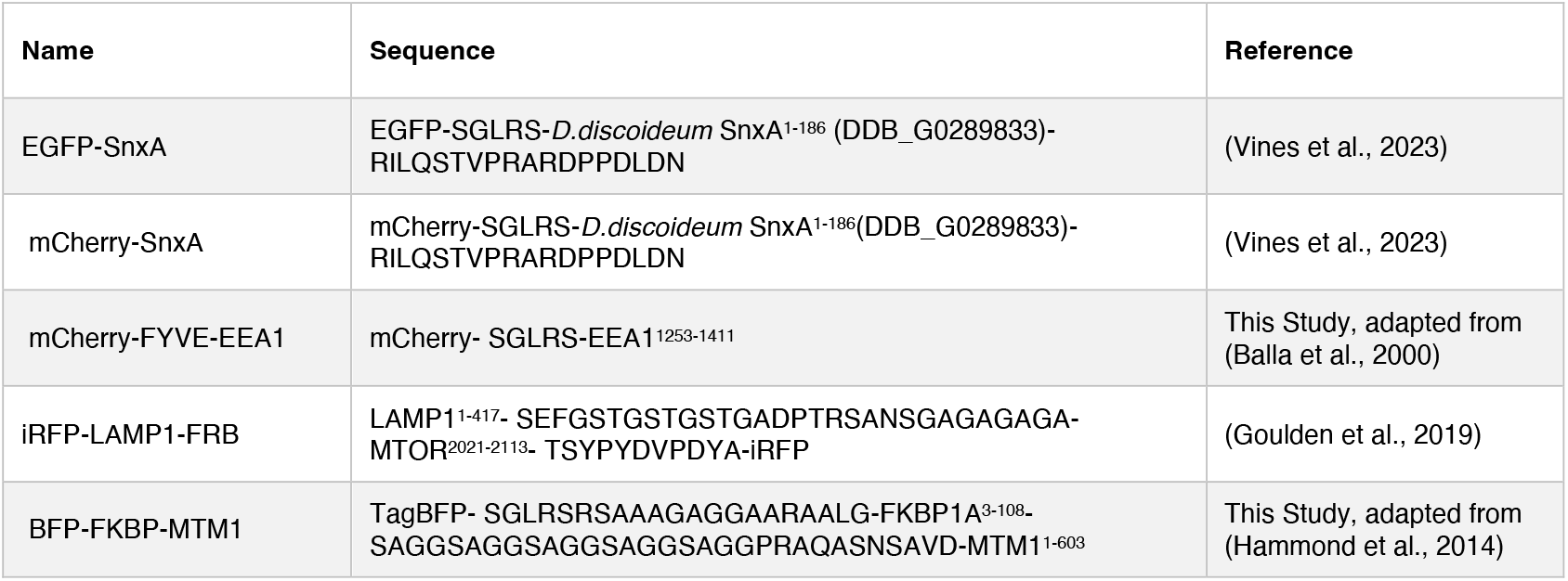
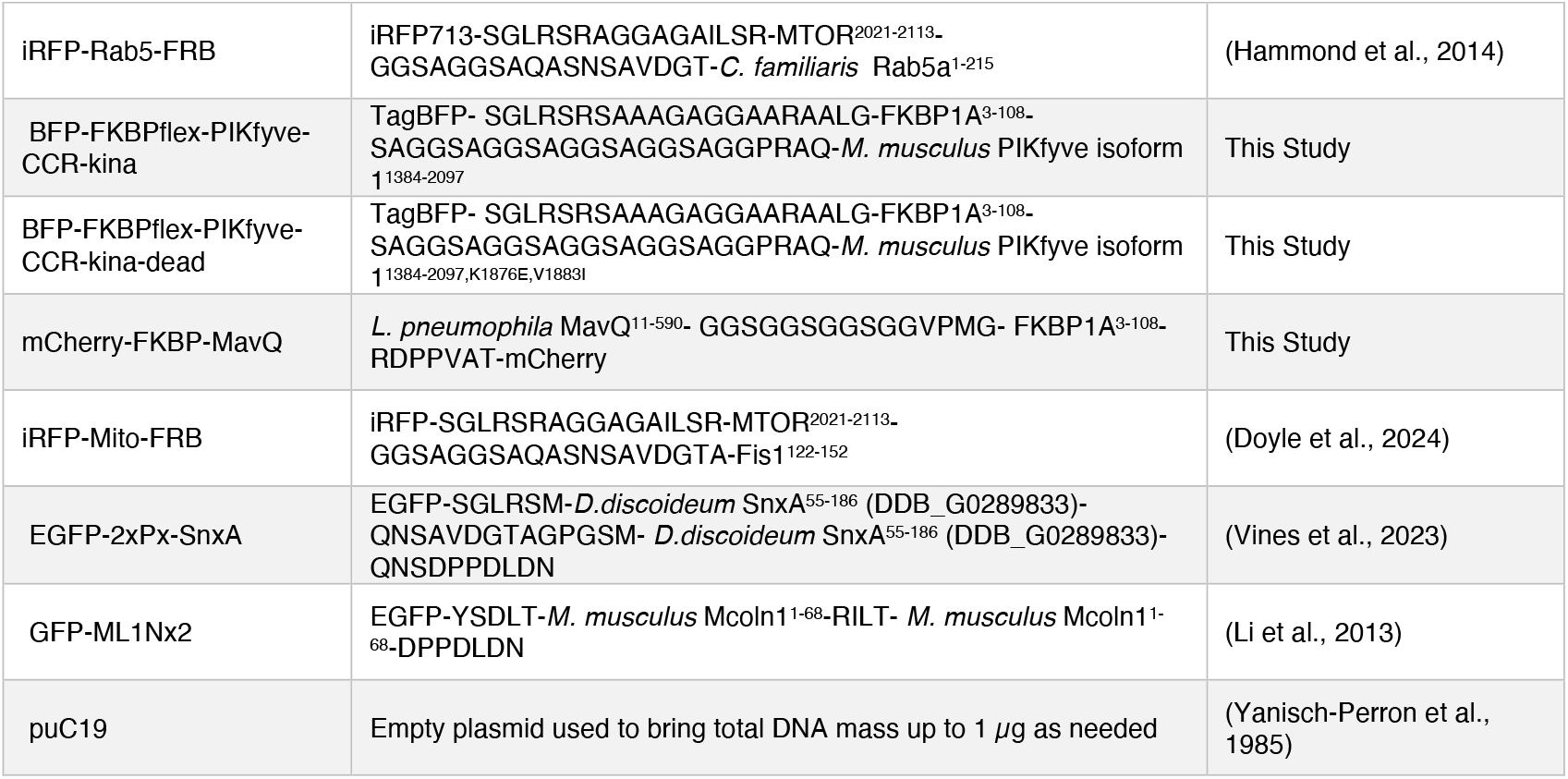
Plasmids used in this study. Numbered residues refer to their position in the full-length protein. All gene are *H. sapiens* unless otherwise specified. Residues written out in 1-letter code indicate linker sequences.

### z-stack Imaging

To image z-stacks of the cells, the confocal microscope was used as described above, with the additional use of the Nikon A1 Piezo Z Drive. The z-stack was set relative to a manual slice set in the middle of the cell. Then, the stack imaging moved from bottom to top in 0.175 µm steps across a range of 10 µm (-5.00 and +5.00), with focus been maintained during time-lapse by the “perfect focus” feedback system. To present and analyze these images, only 11 slices in the middle of the cell were used as an average z-projection, as these slices contained the majority of the SnxA-labelled vesicles.

### Confocal Image Analysis

FIJI was used to complete all image analysis (Schindelin et al., 2012). To quantify the intensity ratios of constructs, a membrane marker was used to create a binary mask: either LAMP1-FRB, dextran, Rab5-FRB or Mito-FRB.

The membrane marker’s fluorescence underwent Gaussian blur filters at 4x, 3x, 2x, or 1x the size of the Airy disc of the fluorophore. The resulting images were then subtracted from the image created by the next smaller filter in order to create wavelets. The wavelets were multiplied together and had a threshold of 3x the standard deviation of the original image applied. A 1 or 2-pixel dilation cycle was used on the wavelets image and the background was subtracted using a threshold of 1.5x the mode of the fluorescence intensity. Cells that moved too much during the time course or were dead/dying as determined by their morphology in the DIC channel were excluded from analysis.

The intensity of the construct within the mask was measured and then divided by its intensity throughout the whole cell (as defined by a manually drawn ROI) to create the given ratio. For further details on analysis, please refer to (Wills et al., 2021) . All quantification was plotted using GraphPad Prism 9 or later.

